# Dysfunctional feedback processing in methamphetamine abuser; evidence from neurophysiological and computational analysis

**DOI:** 10.1101/2022.11.08.515688

**Authors:** Sadegh Ghaderi, Jamal Amanirad, Mohammad Hemami, Reza Khosrowabadi

**Affiliations:** Institute for Cognitive & Brain Sciences, Shahid Beheshti University, Tehran, Iran

**Keywords:** Methamphetamine use disorder, Feedback processing, Reinforcement learning, value sensitivity, electroencephalography, theta power

## Abstract

Methamphetamine use disorder associated with a dysfunctional neural feedback (reward-punishment) processing system and is considered a public health risk. Although several behavioral, computational, and electrocortical studies have explored feedback processing in other groups of individuals, the precise mechanisms of feedback processing dysfunction in methamphetamine use dependent (MUD) individuals remain unclear. Furthermore, our recent knowledge about the underlying feedback-related connectivity patterns and intertwining latent components of behavior with electrocortical signals in MUDs remained quite poor. The present study intended to fill these gaps by exploring the behavioral and electrocortical responses of abstained MUDs during a feedback-based learning paradigm. As mathematical models revealed, MUDs have less sensitivity to distinguishing optimal options (less sensitivity to options value) and learned less from negative feedback, compared with healthy controls. The MUDs also presented smaller medial-frontal theta (5–8 Hz) oscillations in response to negative feedback (300-550 ms post feedback) while other measures responsible for learning including, feedback-related negativity (FRN), parietal-P300, and a flux originated from medial frontal to lateral prefrontal remained intact for them. Further, in contrast to healthy controls, the observed association between feedback sensitivity and medial-frontal theta activity is eliminated in MUDs. We suggested that these results in MUDs may be due to the adverse effect of methamphetamine on the cortico-striatal dopamine circuit, reflected in anterior cingulate cortex (ACC) activity as the best candidate region responsible for efficient behavior adjustment. This study unveils the underlying neural mechanism of feedback processing in individuals with methamphetamine use history and could offer individual therapeutic approaches.

## Introduction

Substance use disorders (SUDs) are associated with loss of control over drug-taking behavior regardless of its negative consequences (R. Smith et al., 2020). Methamphetamine is considered as one of the most widely prevalent illegal narcotics in the world and chronic use of it can alter the feedback processing system of the brain (Lim et al., 2021; R. Smith et al., 2020; N. D. Volkow, Chang, Wang, Fowler, Ding, et al., 2001; Wei et al., 2021; Zhong et al., 2020). Hence, methamphetamine abuse can induce deficits including insensitivity to the negative consequences of its consumption (Lim et al., 2021; N. D. Volkow, Chang, Wang, Fowler, Franceschi, et al., 2001). It seems feedback processing network impairment effects not only limited to real-world behavior, and drug-seeking behavior regardless of its negative outcomes, but also overshadows to computer tasks as other sides of decision making (Kluwe-Schiavon et al., 2020; Potvin et al., 2018). It’s suggested that the abovementioned impairment in methamphetamine use dependent (MUD) individuals in the laboratory setting, failure in learning from negative feedback, is due to improper negative feedback (reinforce) processing (Wei et al., 2018; Zhong et al., 2020; see Kamarajan et al., 2014 for other group of addicts). To strengthen this vision behavioral, mathematical modeling, and neuroimaging studies have been investigated separately but not in the form of an integrated study (Lim et al., 2021; Wei et al., 2018; Zhong et al., 2020).

The various feedback-based learning paradigms are used to uncover the reinforcement learning (RL) process in individuals with altered feedback processing systems including ADHD (Frank, Santamaria, et al., 2007; Pedersen et al., 2017), schizophrenia (Collins et al., 2014; Kirschner & Klein, 2022a), SUDs (Harlé et al., 2015; Lim et al., 2021; Pilhatsch et al., 2020; Wei et al., 2018), and etc. In RL contexts, which are associated with uncertainty in the consequence of the choices, a subject should properly learn from the outcomes of his/her action(Cavanagh et al., 2019a; Maia & Frank, 2011). Briefly optimal decisions are choices that maximize reward and minimize punishment and individuals with a history of methamphetamine abuse have a poor performance in this optimality; they are less accurate compared to healthy in learning from feedback (Dean et al., 2018; Dickinson, 1970; Verdejo-Garcia et al., 2018; Wang et al., 2018; see Potvin et al., 2018 for review). This deficit may be associated with long-lasting effects such as relapse despite the active treatment they received(Konova et al., 2020; Nestor et al., 2011). Therefore, fair amount of knowledge about the RL process in the MUDs helps to comprehend the behavioral mechanisms better and as a result, improves the personalized treatment strategy. In this regard, due to the computational model’s potential to provide quantitative hypotheses at the neuronal and behavioral levels, using them for an optimal description of learning mechanisms is promising and helpful because formulate processes that are not readily available (Collins & Shenhav, 2022; Miletid et al., 2020). Noteworthy, computational model parameters can also have specific cognitive interpretations and are considered cognition descriptors(Pedersen et al., 2017). Assign the value to options, the rate of using prior action-outcome information, and the environment exploration to maximize utility can be mentioned as the most important learning components that are described by the models(Collins & Shenhav, 2022; Miletid et al., 2020; Pedersen et al., 2017). Fitted models to addict’s data compared to healthy individuals revealed differences in some learning-related parameters (Gueguen et al., 2021; Myers et al., 2016, 2017; R. Smith et al., 2020). Modeling study using the Markov decision process and hierarchical Bayesian group analyses has shown that abstained substance abusers, including methamphetamine, have a lower negative learning rate compared to positive learning rate (less ability to learn from negative feedback compared to positive feedback)(R. Smith et al., 2021). This reduced sensitivity to losses/negative feedback in other groups of addicts is also been reported (Ahn et al., 2014; Myers et al., 2017). Another study, which fitting accuracy data of probabilistic learning task performed by active addict patients with the RL model confirmed in addition that addicts have significantly lower negative learning rates and intact reward learning rates, they also been less sensitive to value differences compared with healthy controls (Lim et al., 2021). These abnormalities, inconsistency in choices, and lower negative learning rate are also found in other feedback-based learning contexts, in other groups of addicts (active users :Stout et al., 2004, abstained: Scherbaum et al., 2018, see Gueguen et al., 2021 for review). Neural correlates of feedback-based learning mechanism, focused on feedback processing, have been investigated extensively (Cavanagh et al., 2009, 2010; Duprez et al., 2020; Kamarajan et al., 2012; Khamassi & Quilodran, 2010; C. di B. Luft et al., 2013; Nassar et al., 2019; Oberg et al., 2011; Parvaz et al., 2015; van de Vijver et al., 2011; Zhong et al., 2020;see Cavanagh & Frank, 2014 to overview).

The underlying mechanism of change in brain activities during the feedback processing could give us a better insight to understand the nature of reinforcement learning. In this regard, electrocortical signals including Event-Related Potentials (ERPs) recorded during feedback-dependent tasks typically disclose three correlates of feedback processing: (a) The feedback related negativity (FRN), a negative deflection that is more negative for negative/unfavorable/losses/incorrect feedback compared to positive/favorable/gains/correct and elicited around 200-350 ms in the medial-frontal region, (b) the feedback-related parietal-P300, which is related to later cognitive processing including attention allocated to process outcome valence, emerged around 300-500 ms and is more positive for the positive feedback compared to the negative feedback, (c) an increase in medial-frontal theta (MFT) activity around 200-600 ms after the feedback onset, that induces larger power for negative feedback compared to positive feedback(Cavanagh & Frank, 2014; Glazer et al., 2018; C. D. B. Luft, 2014; Martin et al., 2018; Polich, 2007). Considerable work have attempted to link the EEG signals during feedback-based tasks to feedback processing regions (network) in the frontal cortex (specifically a network that includes the cortico-striatal dopamine circuit) and task performances (Heydari & Holroyd, 2016; Holroyd & Coles, 2002; Kamarajan et al., 2012; C. di B. Luft et al., 2013; Oberg et al., 2011; Wei et al., 2018; Zhong et al., 2020; see Glazer et al., 2018 to review). For example, by considering the direct association between dopaminergic midbrain neurons and prediction error (mismatch between expected and obtained feedback) signals in the Anterior Cingulate Cortex (ACC), negative/unfavorable/losses/incorrect feedback is assumed to cause disinhibition of neurons in the dorsal ACC that provokes the FRN(Holroyd & Coles, 2008; Holroyd & Yeung, 2003). Briefly and in inconsistency, some studies assumed that amplitudes of this valenced-dependent component may encode prediction errors (PE) and modulates by learning (Cavanagh et al., 2010; van de Vijver et al., 2011). Due to the role of ACC, it is thought that MFT is a reflection of PE, associative memory, working memory, and plays a vital role in optimizing subsequent behavior based on received negative feedback. In addition to linking between MFT and options value even in uncertain contexts (Backus et al., 2016; Cavanagh et al., 2010; Cavanagh & Frank, 2014; van de Vijver et al., 2011), this oscillatory activity is also associated with performance monitoring and underlying communication provided for facilitating cognitive control and behavioral adaptation (Cavanagh et al., 2009, 2012; van de Vijver et al., 2011; Weismüller et al., 2019). Learning success and options value can also be related to this oscillation(Cavanagh et al., 2010; Cavanagh & Frank, 2014; C. D. B. Luft, 2014; Martin et al., 2018; Oberg et al., 2011).

Investigation on feedback processing mechanisms in addicts, as individuals with dysfunction in network and related regions, have shown that these patients have abnormalities in feedback processing, specifically negative feedback, compared to healthy controls. In this regard, a previous study has shown that problem gamblers have the same FRN amplitude and smaller MFT in response to negative feedback as compared to healthy controls (Oberg et al., 2011). This insensitivity to negative feedback, mirrored in smaller MFT, has previously been reported for abstinent alcohol addicts (see Kamarajan et al., 2014 to overview). Such results for abstained MUDs are rare and inconsistent(Wei et al., 2018; Zhong et al., 2020; see May et al., 2020 for review). For example, in contrast to Wei and colleagues(Wei et al., 2018), Zhong et al.(Zhong et al, 2020) found that MUDs have reduced FRN amplitude (less negative) compared to controls; they do not assess the frequency measures. Also, Wei and colleagues found increased parietal P300 amplitude for MUDs compared to controls.

Hitherto, due to substance effects on the individual’s feedback processing network, cause non-optimal negative feedback processing and as a result, they do not learn from negative feedback sufficiently (Franken et al., 2007; Kamarajan et al., 2012; Oberg et al., 2011). Other studies have shown that MUD’s are associated with a reduced capacity to apply cognitive control due to the impairment that methamphetamine does to the medial frontal cortex, in particular to the ACC (Kluwe-Schiavon et al., 2020; May et al., 2020). Therefore, these patients are less able in performing tasks subject to learning from feedback and subsequently adjusting behavior based on received errors or negative feedback (May et al., 2020; Wei et al., 2018; Zhong et al., 2020). So, it seems that they continue their mistakes (non-optimal choices) even after receiving the error, which slows down the process of updating the option’s value (May et al., 2020; Wei et al., 2018; Zhong et al., 2020). To previous findings confirmation, previous studies have noted the important role of dopamine receptors in the mesencephalon in preference of learning from positive or negative feedback(Klein, 2008; Poulton & Hester, 2020). Briefly, they demonstrated that poor dopaminergic tone in the mesencephalon, which typically substance-dependent individuals are characterized by, may describe their insensitivity to negative feedback compared to positive feedback(Poulton & Hester, 2020).

The current study was conducted in the favor of growing consensus on the existence of MUDs-related deficits in feedback processing system. In addition to classical behavioral analysis, to uncover the microstructure of feedback processing, we also analyzed both electrocortical and latent components associated to behavioral responses related to learning from feedback during probabilistic learning task. We also supposed this context may imitate MUD’s real-world hyposensitivity to the negative consequences of his/her drug-taking. Taken together, we expect at the behavioral level, the MUDs choose less optimal options than controls and, in turn, this suboptimal selection in MUDs would be due to their fewer tendencies to learn from negative feedbacks that directly influence their valuation system (less sufficient in distinguishing between optimal and non-optimal options). Moreover, at the electrocortical level, ACC-dependent functions including its association with the cognitive control areas are impaired in MUDs and specifically the MUDs have less MFT power in response to negative feedback as compared to controls. Subsequently, the MUDs cannot use negative feedback information adequately to gain sufficient consistency in choices. This insensitivity to punishment in MUDs will reflect in MFT power while responding to negative feedback during the training phase, the MFT power in MUD fails to predict avoidance learning in the test-phase.

## Materials and Methods

### Participants

Two groups were recruited including 21 male right-handed participants with a history of methamphetamine addiction from the Government addiction camp (Tehran, Iran), and 20 healthy controls who were matched in terms of age and gender with the MUDs. For the addict group, the psychiatrist’s diagnosis based on the DSM-IV criteria, self-declaration, and urine test upon entering the compulsory camp confirmed the participant’s addiction. Importantly, to prevent the effects of physical pain and alleviate cognitive disorders caused by methamphetamine use, all MUDs with a history between 21 and 90 days in the camp were included in the study process. The predominance of methamphetamine consumption in addicted individuals, no history of brain damage, neurological disorders, and not taking any psychiatric medication considers as the main inclusion criteria in this study. In addition to the abovementioned screening criteria, the participants were interviewed by a psychologist for cognitive and psychotic’s disorders and three participants were excluded from the study process. For the healthy control group, participants without any history of substance use and any other DSM-IV disorders were recruited through social media. All individuals completed written informed assent before beginning the experiment and they were paid on average 7$ as the collected points at the end of the task. Data were collected in accordance to Helsinki federation protocol and was approved by the ethical board of the university with the ethical code of IR.SBU.REC.1400.008.

### Procedure

After screening and completing Barratt Impulsiveness Scale (BIS) questionnaires which measure impulsivity, individuals performed a task with stimulus-response demand concurrent with EEG recording. Task and all related EEG triggers were conducted in MATLAB in combination with Psychtoolbox. Following presenting task related instructions on the screen 120 trial warm-up provided and some participants information like age and year of abuse, abstain day were requested before tasks. Moreover, based on the 5% nominal test, the accuracy more than 54% during the 600 trials indicates that the individual did not just perform random behavior. Therefore, individuals with accuracy less than this threshold were excluded from the next stages of analysis. However, two addicts whose accuracy was 52% were included in the study because their accuracy (learning success) was acceptable under easier conditions; see table 1 for demographic information (after considering exclusion criteria) of both groups.

**Table 1.** Mean, SD, and statistical results of demographic and behavioral performance.

|  | HC<br>Mean (SD) | MUD<br>Mean (SD) | T | P |
| --- | --- | --- | --- | --- |
| Age | 27.5(3.75) | 28.07(4.3) | -0.3 | 0.74 |
| BIS | 61.6(6.5) | 73.5(10) | -3.7 | 0.001 |
| Year of abuse | - | 5.3(3.5) | - | - |
| Abstain day | - | 57(27) | - | - |
| Training accuracy (%) | 65(9) | 60(7) | 1.6 | 0.1 |
| Training RT (sec) | 1.17(0.25) | 1.21(0.24) | -0.53 | 0.59 |
| AB accuracy (%) | 74(16) | 67(12) | 1.3 | 0.18 |
| CD accuracy (%) | 56(16) | 53(13) | 0.7 | 0.48 |
| EF accuracy (%) | 65(9) | 61(13) | 0.84 | 0.4 |
| AB RT | 1.11(0.26) | 1.19(0.23) | -0.9 | 0.37 |
| CD RT | 1.22(0.24) | 1.28(0.24) | -0.6 | 0.5 |
| EF RT | 1.17(0.26) | 1.17(0.25) | 0.02 | 0.98 |
| Test- Nogo accuracy | 64(21.8) | 58(27) | 0.7 | 0.48 |
| Test- Go accuracy | 81(21.3) | 64(20) | 2.2 | 0.03 |
| Test high Conflict Nogo accuracy | 55(17) | 54(18) | 0.17 | 0.76 |
| Test low Conflict Nogo accuracy | 57(24) | 57(28) | 0.26 | 0.79 |

### Behavioral assessments

Different aspects of impulsivity, including nonplanning, motor, and attentional aspect was measured using the *Barratt Impulsiveness Scale* (BIS) (Patton Jim H, Stanford Matthew S, 1995).The BIS is a 30-item self-completion questionnaire that scored between 0 to 4 and is designed to measure these three different aspects. Higher scores in this valid questionnaire indicate greater impulsivity.

### Experimental task

We used a modified version of the probabilistic learning task (PLT) which comprised 600 trials training phase followed by a subsequent 120 trials testing phase (Fig. 1)(Frank et al., 2004). In each trial of the training phase the participants were presented with three stimulus pairs (Japanese Hiragana character) with a dissimilar probability of receiving “Correct” or “Incorrect” feedback. The stimulus couples (and their probabilities of being rewarded) were according to A/B (70%/30%), C/D (65%/35%), and E/F (60%/40%). As a procedure, all trials initiated with a jittered inter-trial-interval (ITI) between 300 and 500ms. Each stimulus pair were displayed randomly on the screen for a maximum of 3000ms, or until the choice was made by marked related items on the keyboard (Z: marked as left, M: marked as right). The result of button press was presenting “correct” or “incorrect” feedback for 750-ms (50^~^100ms post button press). Also “No Response” was presented when the participants didn’t choose within the 3000ms.

**Figure 1.**
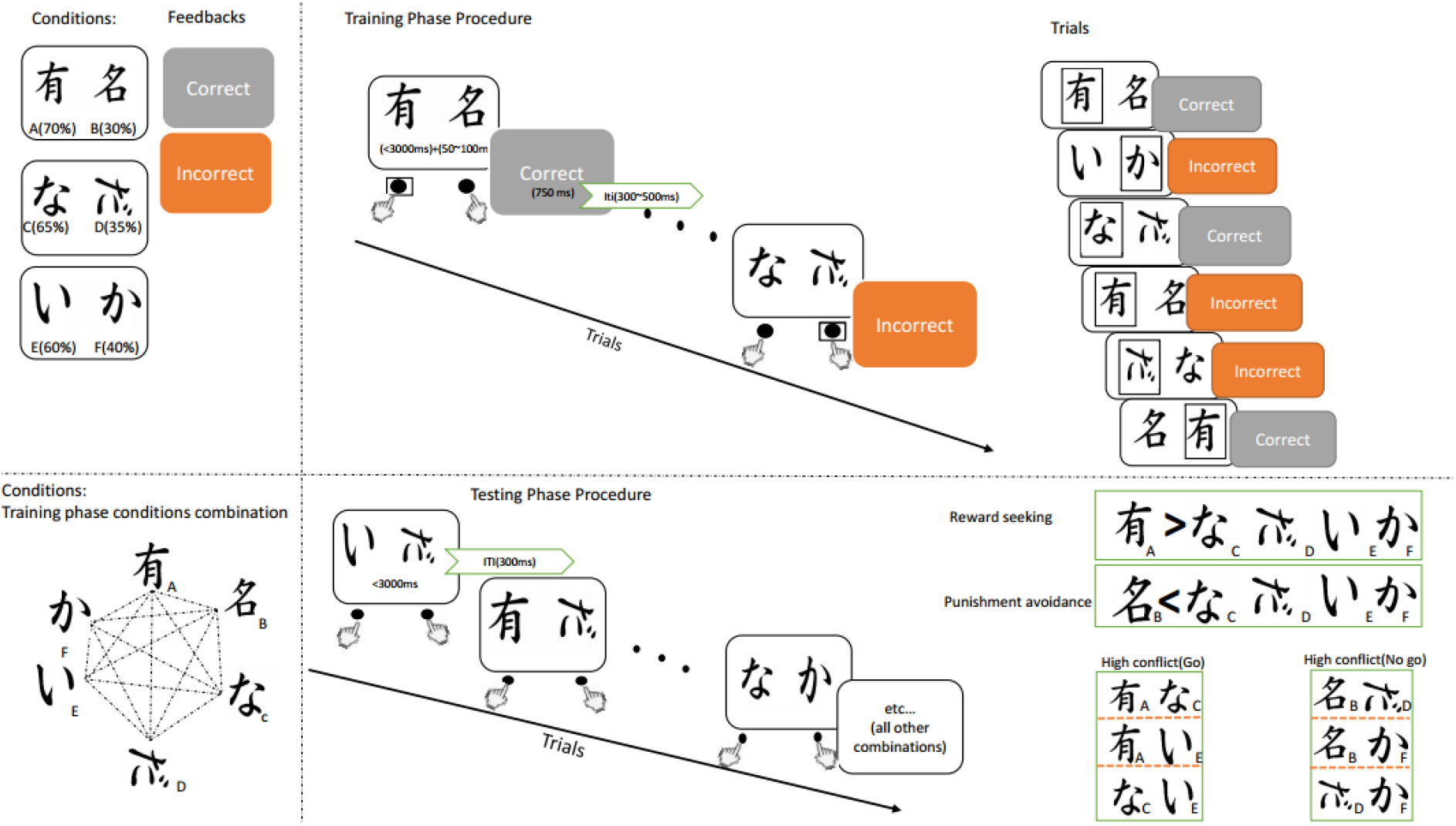
Probabilistic learning task. In each trial of the training phase, individuals have to choose one of the stimuli of the three presented pairs. Each stimulus has a reward/punishment in probable rate and after any selection, feedback (“correct” or “incorrect”) is presented on screen. In the testing phase, all possible combinations of stimulus are presented separately and there is no sign or feedback provided for choosing and individuals should use what they learned from the training phase.

In the course of the testing phase, all possible stimulus combination (e.g., AB, AC, etc.) presented eight times for a maximum of 3000ms and vanished instantaneously after the stimulus selection. Each trial of this phase begins with 300ms ITI. There is no sign or feedback provided in the testing phase and individuals should use what have learned in the training phase to get maximum accuracy for each type of condition. Importantly, there are different interpretations for different combination stats and their corresponding response. Hence, “Go learning” or reward seeking was defined as the accuracy of choosing A in AC, AD, AE, and AF trials. Punishment avoidance or “NoGo learning” was defined as the accuracy of choosing C, D, E, and F in BC, BD, BE, and BF trials. Other conditions which provided in this phase are high/low reinforcement conflict decisions [high conflict NoGo (BD, BF, DF), and low conflict NoGo (BC, BE)]. Accuracy in these combinations is considered as more avoiding B.

### Computational modeling of behavioral data

To describe between trials dynamic of training-phase performance data we fit trial-by-trial sequence of choices for each subject by applying *the Q*-learning reinforcement learning (*Q*-RL) model(Frank, Moustafa, et al., 2007).

*Q* -RL is formalized as follows:

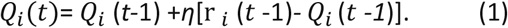

Where subject expectation from the outcome for chosen alternative in trial *t*, *Q_i_*(*t*), is influenced by the difference between the observed (r) and expected outcomes, previous trial expectation, and learning rate parameter (*η*).

Hitherto, this model states learning is driven by, learning rates, and the difference between the observed (r) and expected outcomes. Finally, *Q*-values were fed to softmax logistic function in combination to free parameter (*β*) to produce probabilities (*P*) of choices for each trial:

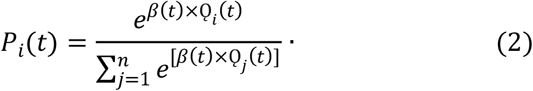

According to the above equation, is the probability of choosing the option with the highest mean reward, and determined behavioral consistency in selection. Generally, *β* is termed as a sensitivity parameter and controls the exploration-exploitation trade-off. Thus, a higher amount of *β* cause the subject to make more deterministic choices (due to being more sensitive to value differences), while it lower values allow the subject to more explore between options.

Practically, the Q-RL model could be included separated learning rate parameters separately for rewards and punishments. Based on model comparison we preferred the model with a separate learning rate; see *Supplementary material A* to the fitting procedure.

### Electrocortical Recording and Processing

Continuous brain electrical activity signals or EEG were acquired using 64-channel active electrodes arranged according to the 10 –20 system and amplified by a g.HIamp amplifier (impedances < 10 kΩ; the online reference was a single channel attached to the right mastoid). The EEG signals were sampled at 512 Hz and were band-pass filters 0.5 to 48 Hz. Offline EEG preprocessing and analyses are performed by the two MATLAB-base toolboxes (in combination with custom scripts): EEGLAB for data preprocessing(Delorme & Makeig, 2004) and FieldTrip custom routines for subsequent analysis(Oostenveld et al., 2011).

#### Preprocessing

After removing the mastoid lead channel from the data structure, the remaining data were epoched around feedback onset (from 300ms before the feedback to 1000ms after the feedback stimulus). Next, large artifacts discarding, bad channel interpolation, and bad epoch rejection (epochs comprising amplitudes above 100μV) were provided automatically and based on visual inspection. Eye blinks artifacts were corrected following independent component analysis. Finally, all EEG channels were re-calculated to an average reference.

#### Event-related potentials (ERPs)

To calculate desired ERPs, data were filtered between 0.5 and 25 Hz and averaged over the trials from −300 to 1000ms after feedback onset. Next, the baseline corrected to the average activity from 300ms before feedback till its onset. The FRN was defined as the mean ERP amplitude between 270 and 300 ms post feedback onset at medial-frontal electrodes (average of electrodes FCz and Fz). The P300 was measured as the mean ERP amplitude averaged over Pz and CPz electrodes between 300 and 450 ms. These time windows were selected based on prior studies and the visual examination of the main grand averaged ERP results(Bedwell et al., 2016; Cavanagh et al., 2019b).

#### Time-frequency calculation

To extract time-varying spectral information of the EEG signal we employed wavelet-based time-frequency decomposition (TFD). First, the signal for each feedback-locked epoch was convolved with a set of varying-cycle complex Morlet wavelets. To regulate the trade-off between temporal and frequency resolution, the number of cycles increased per frequency (frequency range scaled linearly from 2 to 40Hz in the step of one). Importantly, to treat edge effect of filtering we stretched the signal by rounding the maximum trial length up to the next power of 2 using zero padding. Second, for each epoch, the feedback-locked estimates of power elicited from the resulting complex signal. Next, these single-trial TFD averaged across trials for each participant. Finally, to normalize TFD per channel we divided spectral power values in each frequency by its baseline value (−300 to −200ms pre-feedback onset). This consideration, allowed us to compare direct effects of feedback across frequency bands. According to previous studies with other groups of the subject, we consider averaged power over FCz and Fz as the electrode of interest which provide an estimation about the medial-frontal activity(Kirschner & Klein, 2022b; C. di B. Luft et al., 2013).

#### Functional connectivity analysis

To calculate long-range cortical interaction between electrodes of interest, we computed inter channels phase synchrony (ICPS) and averaged over trials. ICPS measures the consistency of phase angles between two electrodes (or regions) over time-frequency points(Reinhart & Nguyen, 2019; van de Vijver et al., 2011; Watts et al., 2018). ICPS is defined as follows:

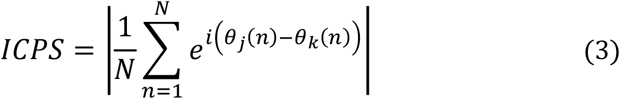

Where *N* is the number of trials for each time-frequency point, is the complex operator, *n* is trial, and *θ_j_* and *θ_k_* are the instantaneous phase values of electrodes and, respectively. Here, to be able to focus on the effect of the feedback on phase coupling, we calculate the percentage changes of ICPS relative to its baseline values (−300 to −200ms pre-feedback onset).

#### Effective connectivity analysis

To provide information about the directionality of long-range connectivity between electrodes/region of interest, we compute PSI. This measure estimate the direction of information flow between two source (EEG channel signal) according to the slope of the phase of their cross-spectrum and is also insensitive to volume conduction artifacts (Nolte et al., 2008). For each couple of electrodes/region the sign of PSI indicates evidence for the role of each electrode/region, a positive value provides evidence an electrode/region acts as a source, and negative values indicate that an electrode/region acts as a sink. Importantly, PSI was calculated over the frequency band of interest and averaged across feedback-locked epochs for each participant and electrode pairs of interest. The frequency band and temporal bin of interest were determined by the ICPS results(see Reinhart & Nguyen, 2019 for inspiration).

### Statistical analysis

To assess trial effect, we fit two separate hierarchical regression models: a logistic regression on accuracy and a linear regression on RTs (see *Supplementary material* B to details). Behavioral task performances model, *Q*-RL results, were analyzed by hierarchal Bayesian inference. This method allowed us to assess the between and within-group difference using posterior distribution of separated parameter based on directional Bayes factors(Wetzels & Wagenmakers, 2012). Electrophysiological responses to feedback were compared using a 2×2 repeated measures ANOVA via a within-participants factor of valence (correct versus incorrect) and the between-participants factor of group (healthy control (HC) versus MUD); then any observing interactions followed by paired and independent *t-*tests.

Next, to investigate correlation between of electrocortical measures with behavioral task performances, we applied time course Spearman’s test. In this approach, if any correlation remained significant for more than 50ms we interpreted it as significance range. Related Statistical analyses were implemented with R-studio, Matlab and PyStan 2.19 custom routines. All data are available on the Open Science Framework (https://osf.io/bercx/) but analysis script would be available upon reasonable request to corresponding author.

## Results

### Behavioral results

By applying exclusion criteria’s, 31 participants (HC =17, MUD=14) data included to main analysis procedure (table 1). Accuracy (optimal choices) was coded logically, consider 1 if the options with the higher probability of reward were selected (e.g., A is selected over B). Mean optimal choices of included participants for healthy control’s and MUD’s were 0.68(SD=0.09) and 0.62(SD=0.07), respectively. Moreover, average training phase optimal choice shift through the course of the task (bin of trials) interpreted as a learning effect (Fig. 2a). It seems, compared to HCs, the MUDs performances were less affected by learning. Such, HCs mean optimal responses varied as bins, from 0.56(the first bin) to 0.75 (last bin) and from 0.53(the first bin) to 0.65(last bin) for the MUDs group. Moreover, consistent with the accuracy-response time (RT) trade-off, mean RTs corresponding to the group also become faster as the task progresses (Fig. 2b). Also, these results confirm by regression analysis while the logistic regression model on accuracy and linear logistic regression model on RT suggest that the trial affected accuracy and RT. In nutshell, the 95% BCI was higher than 0 for the mean group trial coefficient in logistic model (for HCs: 0.04 to 0.06, *mean* = 0.055; for MUDs: 0.026 to 0.045, *mean* = 0.035; BF>100) and lower than 0 for the trial coefficient in linear model only for HCs (−0.046 to −0.038, *mean* = −0.04; for MUDs: 0.001 to 0.009, *mean* = 0.005; BF>100). Statistically, no strong differences were found between MUDs and HCs in direct learning-related performance measures neither in the training phase nor test phase (see table 1).

**Figure 2.**
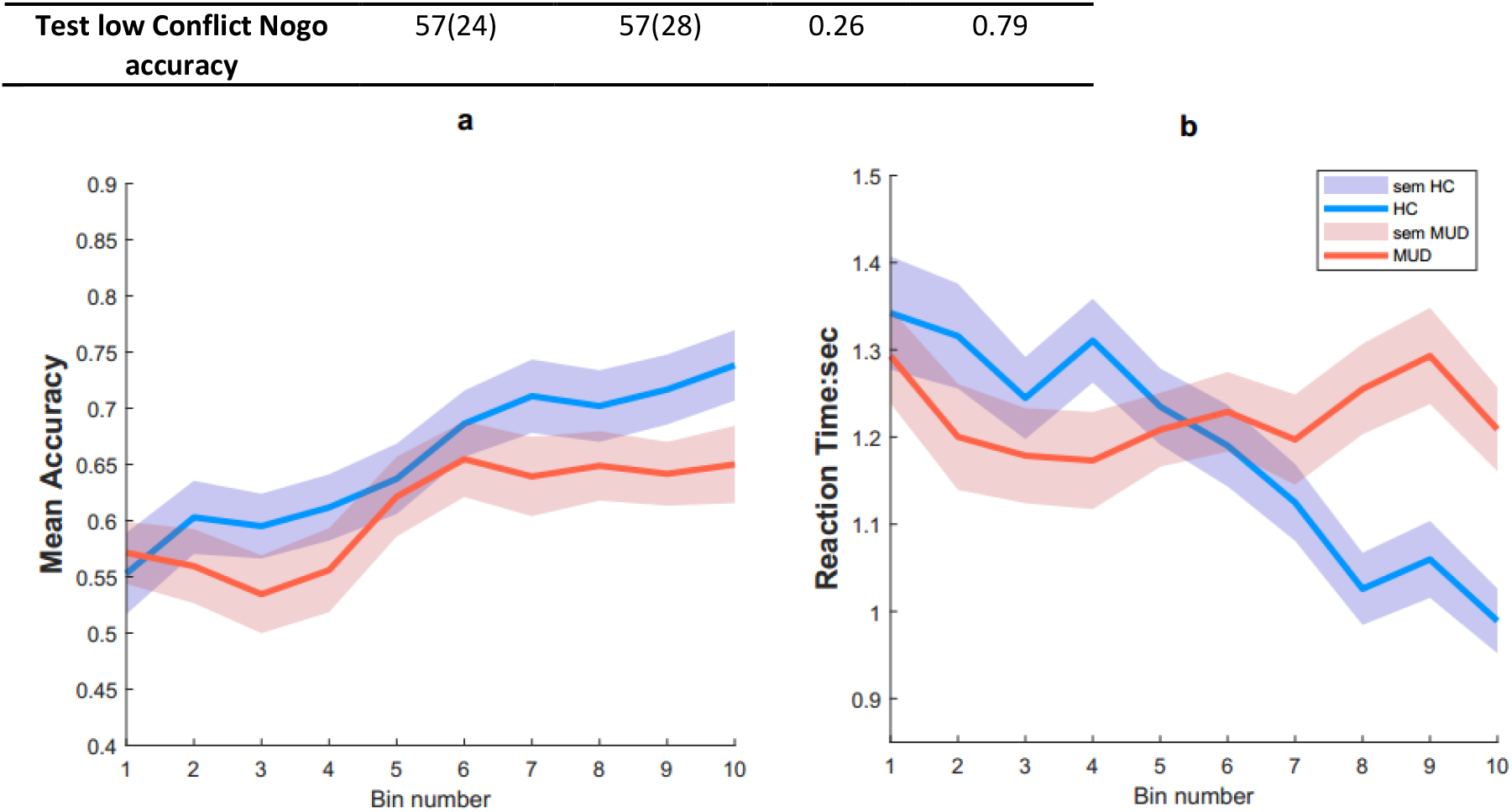
Mean of accuracy (a) and Response time (b) alter by learning across the group (blue: HC, red= MUD). Thick lines characterize the mean across the experimental task bin (every 60 trials =1 Bin) and participants and the shaded areas indicate the standard error of the mean (sem).

### Effect addiction on computational mechanisms of learning

By benefits which provided through model comparison and other models related evidence we went through the rest of the steps by the model with dual learning rates, see *Supplementary material A* (table1). Conducted analyses to interpret between groups differences, by the Bayesian framework, revealed the MUDs underwent less value sensitivity in responses. Moreover, relative to healthy controls, addicts have less negative but same positive learning rates, addressed in Fig. 3; also see *Supplementary material A* (table 2) for more insight.

**Figure 3.**
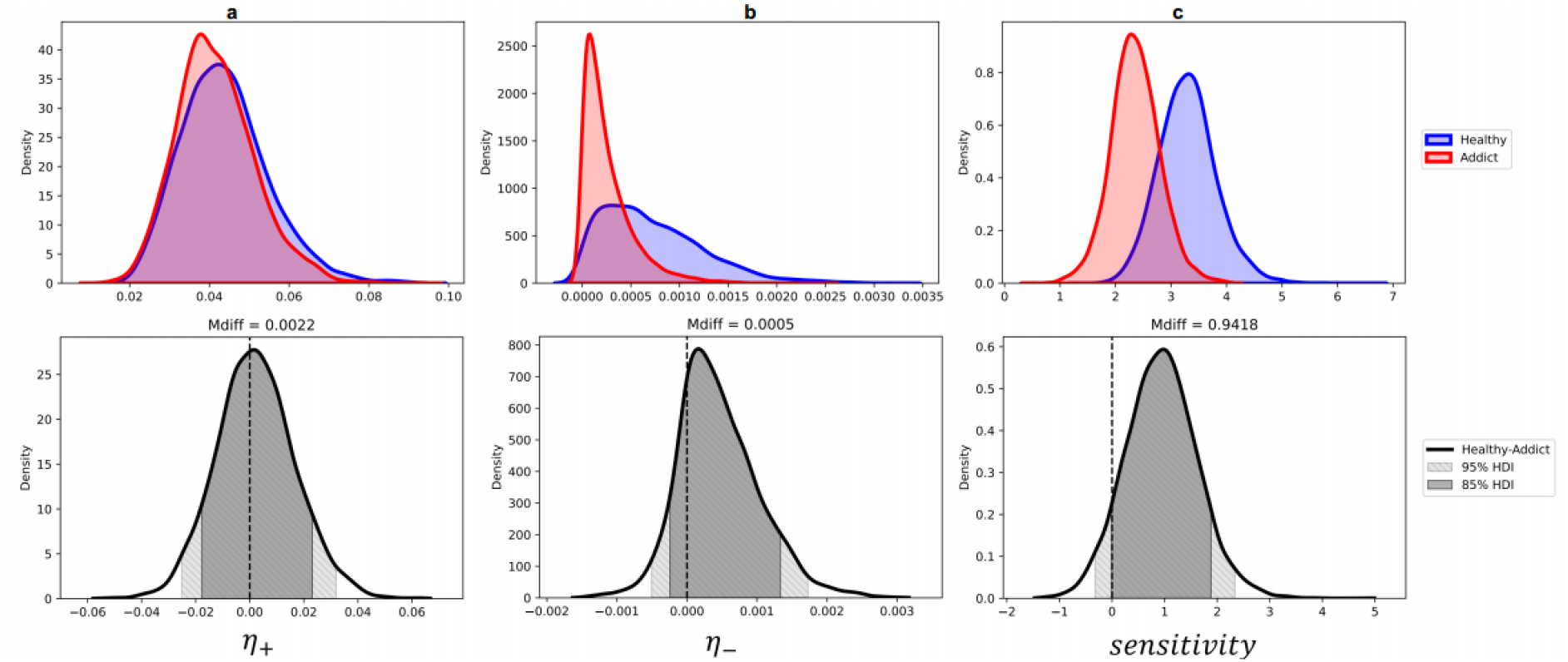
Results for RL model with separated learning rate. Upper panel: Posterior distributions of parameters for HC group (blue) and MUD group (red) for positive (a), negative learning rate (b), and options value sensitivity (c). Lower panel: Mean posterior distributions of group differences per parameter (HC group- MUD group, Mdiff: Mean_HC_- Mean_MUD_). Note: 85% (95%) highest density intervals (HDI) are illustrated by dark gray (light gray) shadowed areas in lower distributions, see *Supplementary material* A to more details including posterior predictive checks (PPC).

### Electrocortical analysis of feedback

#### *ERP* results

We found a negative discharge on ERPs with frontocentral topography which elicited around 270-300ms following negative feedback relative to positive feedback (Fig. 4). The component with these features interpreted as an FRN. A mixed ANOVA (2[feedback] ×2[group]) on the mean FRN amplitude disclosed significant effects of feedback (F _(1, 29)_ =50.16, *P*<0.001, partial **η**^2^=0.1). No other effect of interest, including group and interactions observed (all F<1, *P*>0.1). These findings indicated the FRN was greater (more negative) for incorrect than correct feedbacks (*t*_(30)_=− 7.25, *P*<0.001), while it appeared similar in both groups. Similar examination conducted for parietal P300 unveiled main significant effect of feedback (F _(1, 29)_ =28.15, *P*<0.001, partial **η**^2^=0.03), but no effect for group nor interaction (all F<1, p>0.1). Again, these results estimated that the amplitude of the parietal P300 was greater for correct feedback than for incorrect feedback.

**Figure 4.**
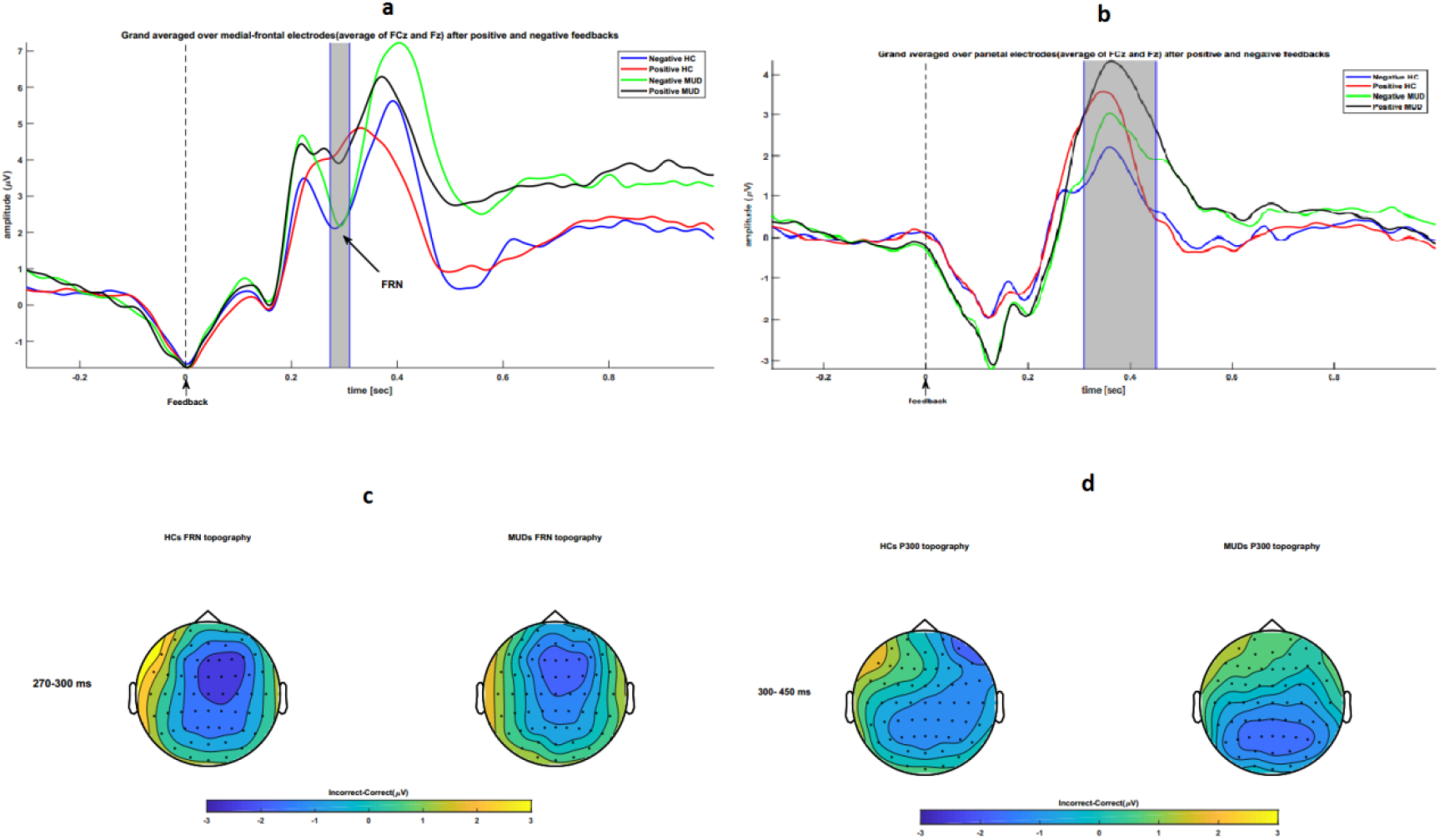
ERPs results for HCs and MUDs following negative and positive feedback. **a**, medial frontal electrodes (averaged over the FCz and Fz) waveforms for the MUD group and HC group in response to negative and positive feedback. **b**, waveforms in parietal electrodes (averaged over the Pz and CPz) for HCs and MUDs following positive and negative feedback. **c**, topographic maps of the FRN during 270 to 300 ms following feedback onset for the differences between negative and positive feedback for HCs (left) and MUDs (right)**. d**, The same subtraction among negative and positive feedback for the parietal P300 time window (300–450ms) illustrated.

#### Time-frequency analysis

Again, a mixed ANOVA on theta relative power averaged over medial-frontal sites (FCz and Fz), from 300 to 550ms following feedback onset, disclosed a significant main effect of feedback (F _(1, 29)_ =5.94, *P*=0.021, partial **η**^2^=0.07), as well as learning group alone(F _(1, 29)_ =8.98, *P*=0.006, partial **η**^2^=0.16), but not feedback-by-group interaction (F<1, P>0.1). Paired *t*-test revealed that the HCs theta power after negative feedback was significantly more than positive feedback (*t*_(16)_=2.21, *P*=0.04); this significantly not observed for MUDs (*t*_(13)_=2.1, *P*=0.055). Moreover, as compared to positive feedback theta power, independent *t*-test revealed that the groups of interest only differed in their theta power in response to negative feedback (*for incorrect feedback: t*_(29)_=3.29, *P*=0.002 while *for correct feedback: t*_(29)_<2, *P*>0.1). These results indicated, as compared with reward feedback, punishment feedback coincided with greater theta band (5–8Hz) power in both groups (Fig. 5a). Also, the relative power enhancement of these rhythms after negative feedback (Fig. 5b), distributed in the medial-frontal part of the scalp (Fig. 5c), is more distinguishable in HCs compared to MUDs.

**Figure 5.**
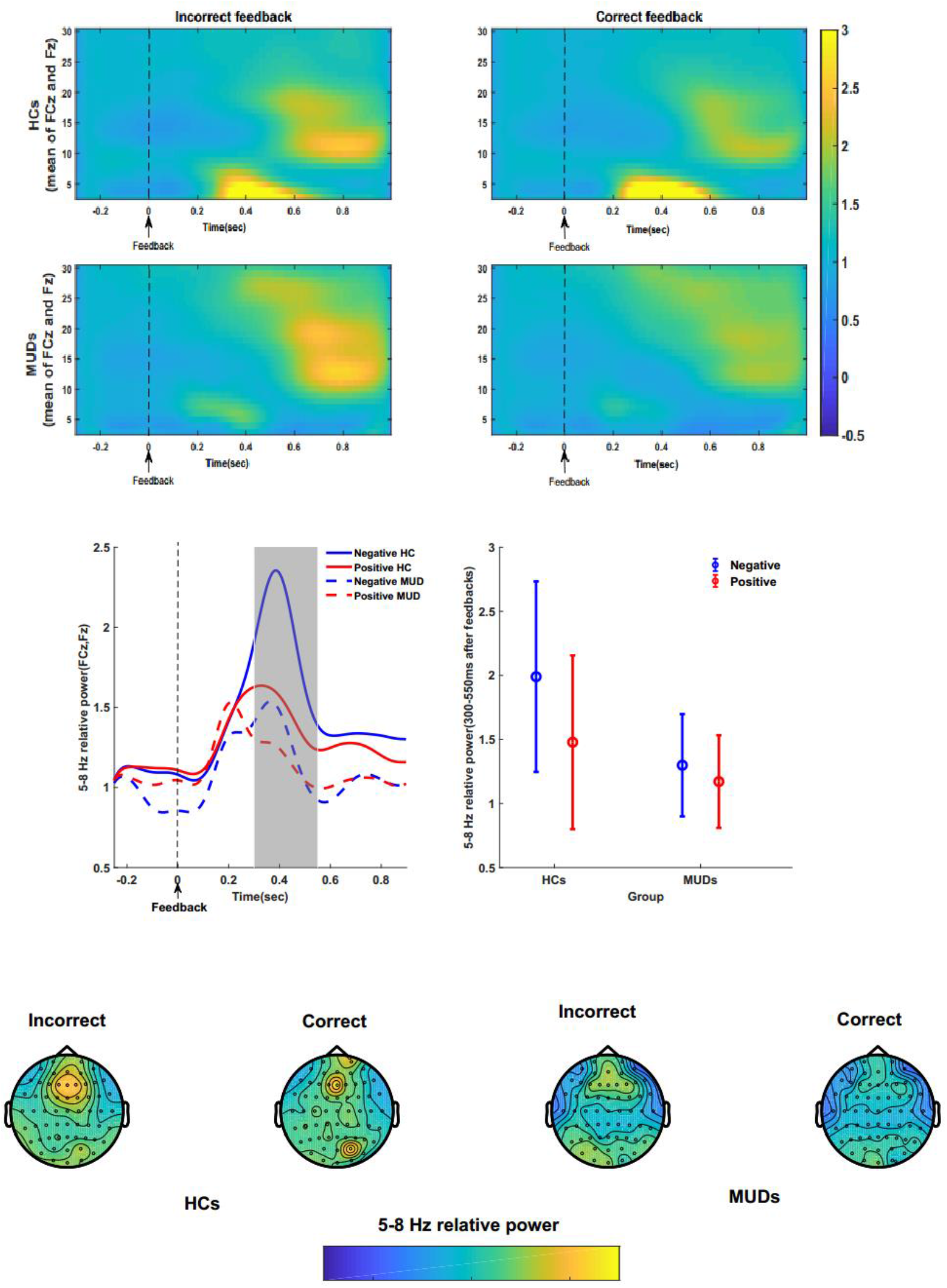
Time-frequency decompositions results. ***Upper panel***, TFDs of EEG spectral power in response to negative (left) and positive feedback for HCs (first row) and MUDs (second row). ***Center panel***, The time course of theta power averaged over FCz and Fz for HCs (solid) and MUDs (dashed) in response to negative (blue) and positive (red) feedback illustrated (left figure). Also, in the right side of central panel the mean and error bar of theta power averaged over FCz and Fz electrodes for the highlighted time window (300-550 ms) on the left side figure is illustrated. ***Bottom panel***, Distribution maps of theta relative power in HC (left side) and MUD (right side) groups during(300-550 ms) negative and positive feedback are shown.

#### Functional connectivity findings

To assess the consistency of low theta (4-6 Hz) phase angles between medial-frontal and right-prefrontal (RPFC) electrodes across groups and feedbacks, we calculated the degree of ICPS percent changes following feedbacks relative to pre-feedback between medial-frontal (averaged over FCz and Fz) and F6 electrodes (Fig. 6a). As compared to positive feedback, we observed more ICPS percent changes for negative feedback between 300 to 550ms post-feedback (see Fig. 6b for waveform). Statistical analyses for (FCz, Fz)-F6 electrodes on this time window revealed the significant effect of feedback (*F*_(1,29)_ =6.78, *P*=0.01, partial **η**^2^=0.038), but no other effect of interest including group and interactions observed (all F<1, *P*>0.1). Taken together, these results indicated that, independent of group effect, the ICPS changes were more significant for negative compared to positive feedback (*t_30_*=2.26, *P*=0.01).

**Figure 6.**
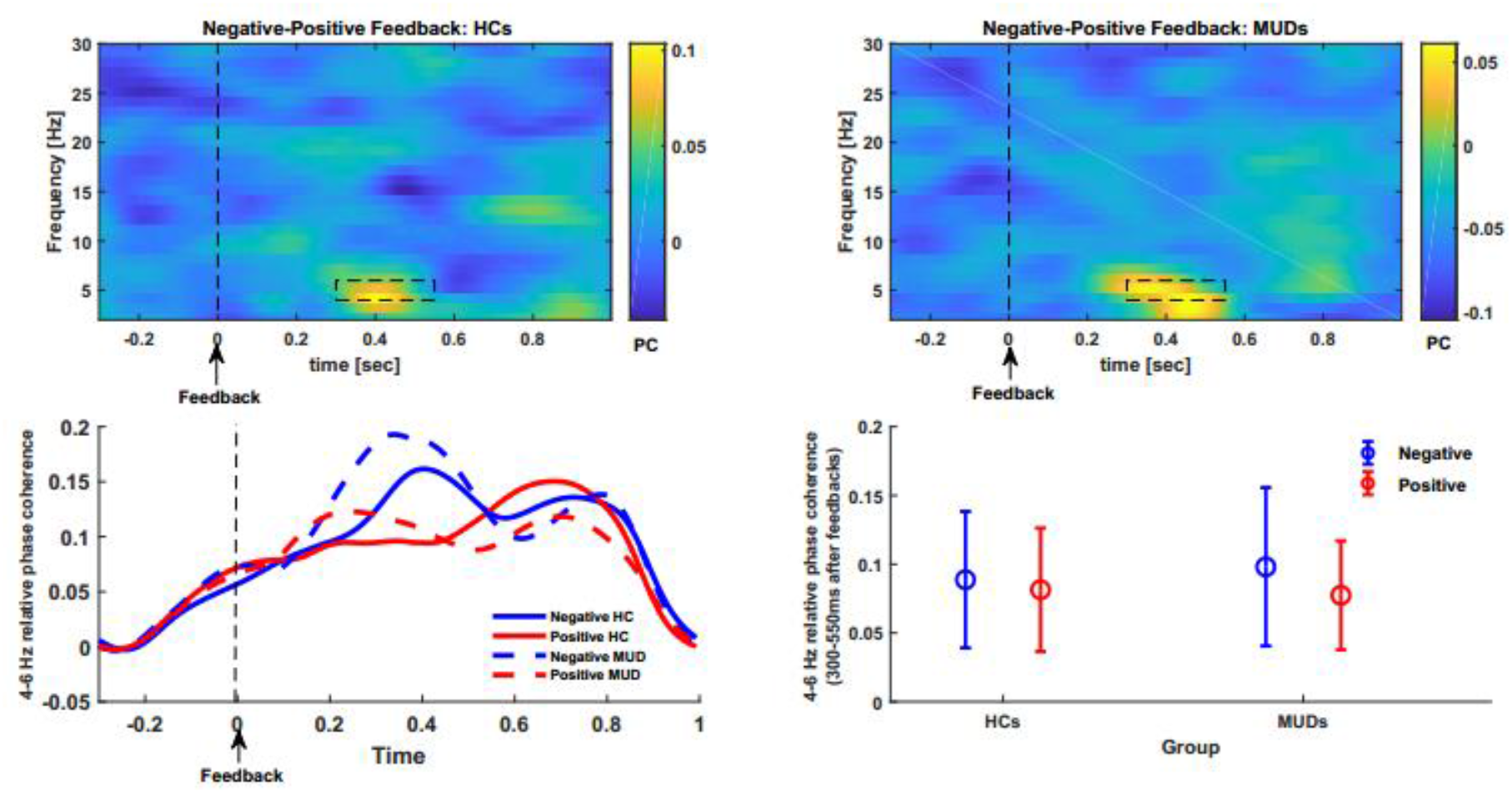
Phase synchrony results. ***Upper panel***, Phase coherency among medial-frontal and RPFC after negative relative to positive feedback for HCs (left) and MUDs (right). ***Lower panel***, The relative feedback-locked theta (4-6) phase synchrony time course for HCs (solid lines) and MUDs (dashed lines) for negative (blue) and positive (red) feedback are illustrated (left figure). The right figures of this panel illustrate the mean and error bar of relative theta-phase synchrony between medial-frontal and RPFC regions for the time window (300-550 ms) of the left figure.

#### Effective connectivity results

Here, the PSI was calculated from 4 to 6 Hz in step of 0.5 Hz and averaged across feedback-locked epochs for each participant and electrode pairs of interest (averaged over FCz and Fz -F6). Temporal bin of interest was 300 to 550ms post feedback onset. We determined this time-frequency ranges based on the corresponded range that showed a significant difference in ICPS changes. Next, to increase the accuracy in detecting PSI we consider ±0.5 HZ window around peaking frequency (5 Hz) as interested frequency band (Fig. 7a). For this time-frequency range we find significant effect of feedback (*F*_(1,29)_ =6.66, *P*=0.015, partial **η**^2^=0.068),but neither for group(*F*_(1,29)_<1, *P*>0.1), nor an interaction (*F*_(1,29)_ <1, *P*>0.1). These findings revealed that independent to the learning group, medial-frontal electrodes provide a leading role in 4 to 6 Hz (by peaking around 5 Hz) connectivity with F6 only after negative feedback compared to positive feedback (*t_30_*=2.4, *P*=0.022), see Fig. 7b.

**Figure 7.**
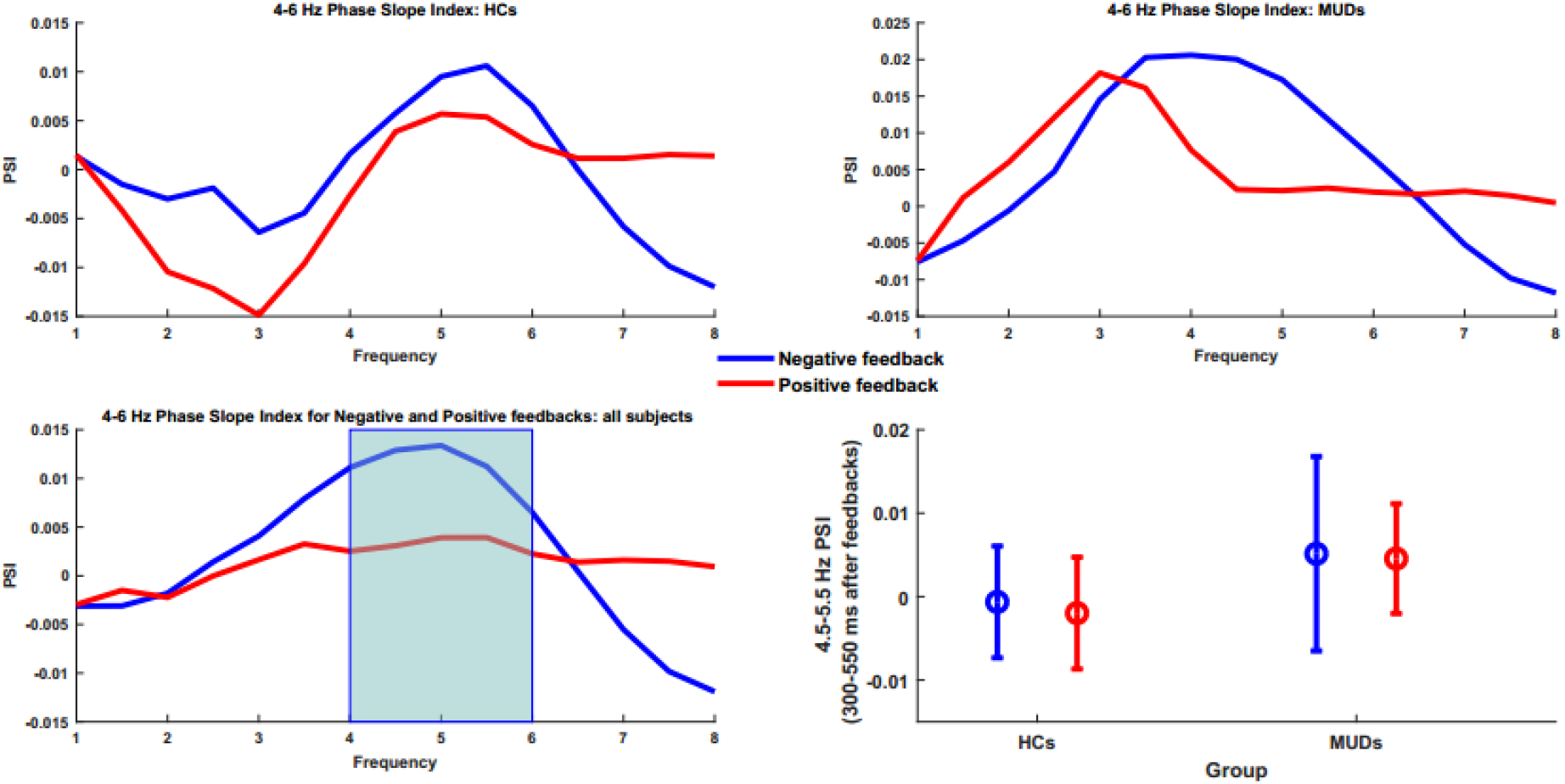
Medial frontal to right prefrontal directionality results. ***Upper panel***, the PSI from 300 to 550 ms following feedback in theta range (4-6 Hz) after negative (blue) and positive (red) feedback for HCs (left) and MUDs (right) is demonstrated. ***Lower panel***, On the left side the PSI across all subjects for negative (blue) and positive feedback (red) which peaked around 5 Hz is demonstrated. On the right side, the mean and error bar of PSI for the time window shadowed (300-550 ms) on the left side is illustrated.

#### Brain-behavior linkage

For the training phase, correlation examination between electrophysiological measures and behavioral task performances including model parameter shows that only theta power correlated with behavioral task performance (Fig. 8a): the higher sensitivity, the higher medial-frontal theta power (5-8 Hz) following negative feedbacks (across all subjects: rho=0.37, *P*=0.04; HCs: rho=0.5, P=0.04; MUDs: rho<0.1, P>0.1). This effect estimated that individuals who had larger theta synchronization in response to negative feedback are more sensitive to value differences hence consistency in behavior. Next, we applied the time course Spearman and demonstrated these oscillatory activities are significantly correlated with sensitivity parameters at the individual level and in narrower intervals beginning around 300 ms and peaking around 400 ms following feedback onset (Fig. 8b). Significantly, group analysis revealed this association only presented in healthy participants; again, began around 300 ms and peaking around 400 ms following feedback (Fig. 8c). For the test phase: at the individual level medial-frontal 5-8Hz power was not associated to avoidance learning neither at the group level (rho=0.28, *P*=0.11) nor the individual level (for HCs: rho=0.37, *P*>0.1, for MUDs: rho=0.01, P >0.1).

**Figure 8.**
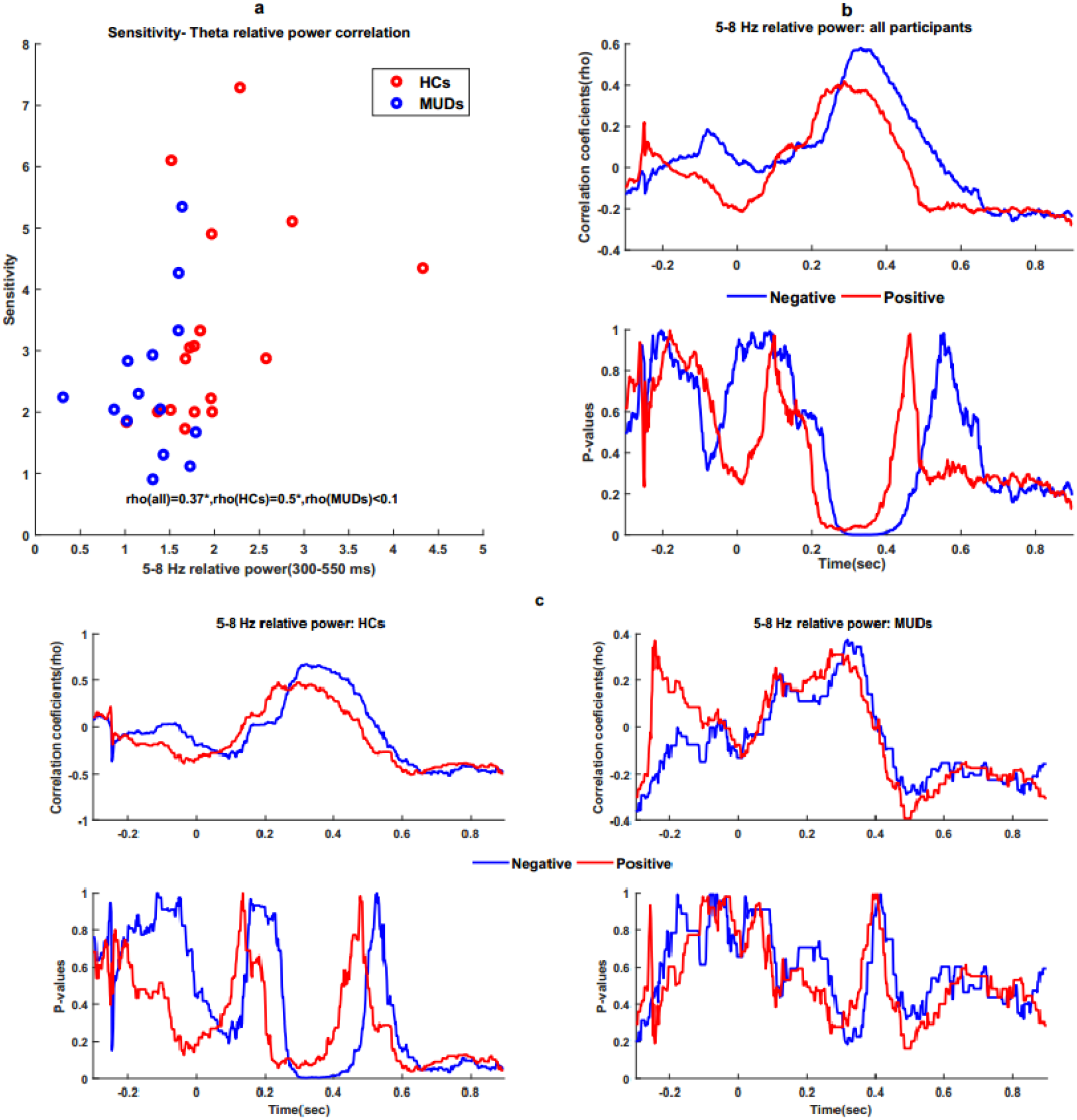
Correlation between theta power and task performance component. ***a***, Scatterplot for the correlation between options value sensitivity and theta (5-8 Hz) relative power from 300 to 550 ms after negative feedback for the HCs (red) and MUDs(blue).***b***, Time-dependent changes of the correlations between theta relative power after negative (blue) and positive (red) feedback and options value sensitivity. The correlation coefficients (top panel) and corresponding *p* values (bottom panel) are illustrated. **c**, Time course of the correlations between theta relative power after negative (blue) and positive (red) feedback and options value sensitivity for HCs (left column) and MUDs (right column); the bottom panels indicate the respective *p* values for HCs(left) and MUDs(right) for negative (blue) and positive (red) feedback.

## Discussion

In this study, we assessed behavioral, computational, and neural characteristics of the feedback-based learning in abstinent methamphetamine abusers and healthy peers through implementing the simple prediction error task. Considering phenotype of addiction(Verdejo-Garcia et al., 2018; Nora D. Volkow & Muenke, 2012), and the deficit in the feedback processing network caused by methamphetamine(Groman et al., 2018; Shukla & Vincent, 2021; N. D. Volkow, Chang, Wang, Fowler, Ding, et al., 2001), we hypothesized that the abstained addicts have malformed learning-related patterns. This malfunctioning is reflected in behavioral, computational, and neurophysiological levels compared to the healthy controls. Subsequently, to test our hypothesis, we examined the separable effects of methamphetamine consumption in three main domains: (A) behavioral performance, (B) performance of the related computational model, (C) electro-cortical responses correlate of behaviour using ERP analysis, investigation of neural oscillations, and pattern of functional connectivity. In addition, association between behavioral performance and neural activities were also computed. Although there have been a few ERP and computational studies on feedback processing dysfunction in methamphetamine use disorders, this is the first frequency-computational study to report feedback processing dysfunction in methamphetamine use disorders.

### Behavioral performance and its latent components

Preliminary learning-related behavioral analyses indicated that HCs and MUDs have similar performance (table 1). Since learning is a time-dependent function, we assessed trial effects by fitting regression models on participants’ accuracy and RTs. We found a significant inferiority in benefits from trials in MUDs compared to HCs. A more detailed examination using the Q-RL model on between trials dynamics indicated that both groups have similar positive learning rate (***η***_+_) while learning rates for negative feedbacks (***η*_−_**) are slightly greater in HCs. This difference suggests that HCs learn faster from negative feedback as compared to MUDs (supplementary material, table 1). It’s also explained why methamphetamine-addicted individuals often fail to learn from the negative consequence of their experiences. Furthermore, our analyses revealed that the HCs were more sensitive to value differences (*β*) as compared to the MUDs. This group difference indicated that HCs benefited more from deterministic choices. Hence, the HCs have more consistency in behavior and could gain optimality in responses (Daw et al., 2006; Pedersen et al., 2017; Sutton & Barto, 1998).

Similar findings have been reported using other learning paradigms and in other groups of addicts. For instance, consistency in choice and influence of negative feedback were found to be lower in active cocaine-dependent individuals in Iowa Gambling Task as a four-armed bandit problem(Stout et al., 2004). The findings were also consistent with those of Scherbaum et al. who compared HCs versus abstained heroin addicts and showed that the addicts individuals have less sensitivity in distinguishing optimal from nonoptimal options (Scherbaum et al., 2018). This unstable response rule was also observed in the reward and punishment learning context for other groups of addicts (Myers et al., 2016). Also in a recent study, using three-armed bandit task and by applying the active inference model, Smith et al. concluded that abstained substance addicts have more learning rate for rewards and a lower learning rate for losses than HCs(R. Smith et al., 2020). The minor differences in the results of these studies specifically for results of positive learning rates could be due to the context, substance use characteristics, and individual covariates. Taken together, our results including observed hyposensitivity for losses and lower options value sensitivity support previous works in SUDs. These works demonstrated that (i) abstained heroin addicts have insensitivity to negative feedback while performing the Iowa Gambling Task. It should be mentioned that this work was performed in combination with the hybrid model of reinforcement learning (Ahn et al., 2014). Similar finding has been reported for for other drug abusers (Haines et al., 2018); (ii) the active SUDs have a lower negative learning rate, similar reward learning rate, and a lower sensitivity to value difference while performing a probabilistic reinforcement learning task (Lim et al., 2021).

Our results also reconciled with the previously discussed dopaminergic system findings. the destructive effect of methamphetamine consumption on different brain regions including the midbrain has been reported in previous studies (Frank & O’Reilly, 2006; Klein et al., 2008; Lim et al., 2021; N. D. Volkow, Chang, Wang, Fowler, Ding, et al., 2001). We suppose this insensitivity to punishment can be addressed by down regulation of striatal D2 receptors and dopaminergic neurotransmission. According to the opponent actor learning model, this disruption can reduce the options value sensitivity (Collins & Frank, 2014; Nagano-Saito et al., 2012; Robertson et al., 2016; see Shukla & Vincent, 2021 to review); Moreover, value sensitivity decreasing in the MUDs can be due to disruption effect on options value updating network including both medial prefrontal cortex (dorsal/ventral rote) and frontostriatal connectivity(Bartra et al., 2013; London et al., 2014; Lopez-Gamundi et al., 2021). Also, poor MUDs working memory (Sabrini et al., 2019) can be another reason underpinning addict individuals’ infirmity in distinguishing optimal options. To reconcile with working memory deficit, recent studies have assumed that the option values may be retained in the working memory. So, it can be discussed that less capacity in working memory may lead to a weak strategy for distinguishing between the options, hence, a lower sensitivity to options value may achieved (Collins et al., 2017; see Yoo & Collins, 2022 to overview).

Finally, our first hypothesis supported while our modeling results provide evidence that estimated methamphetamine abusers are determined by aberrant latent learning mechanisms. Moreover, these results strengthen the view that MUD patients may have a dysfunctional neural mechanism associated with these mechanisms, specifically feedback processing.

### Electrocortical signals in service to learning: from ERPs to connectivity

Converging to previous findings, our ERP results for the FRN component revealed that negative feedback elicited more negative power relative to positive feedback (Glazer et al., 2018; C. di B. Luft et al., 2013; Rawls et al., 2020; Zhao et al., 2017; see Glazer et al., 2018 for review). In contrast to study performed by Zhong et al. (Zhong et al., 2020), our results did not reveal any differential FRN amplitude in healthy individuals as compared to abstained methamphetamine users. It seems the covert reason for observing this discrepancy is the nature of the conducted tasks, the Balloon analog risk task (BART) in Zhong’s study examined risky decisions, while the PLT in our study examined the response outcome contingency. Our between-group results are also contrary to the findings of Wei and colleagues who found that female methamphetamine users had more negative FRN after feedback in a simple gambling task as compared to healthy controls (Wei et al., 2018). However, it is difficult to reconcile their results with the reported ACC hyposensitivity finding which claimed the ACC is less activated in regulating MUDs’ responses to punishment as compared to HCs (Stewart et al., 2014; see May et al., 2020 for review; also see Zhao et al., 2017 for results in another group of addicts). In addition, our results were consistent with the previous findings on other group of individuals with learning disability including autism, and healthy non-learners that pointed out to intact FRN despite observing behavioral malformed patterns (for autism: Hüpen et al., 2016; for healthy non-learners: C. di B. Luft et al., 2013;also see Martin et al., 2018 for overview). Moreover, we found parietal P300 was larger (more positive) for positive feedback than negative feedback in both groups. A previous study suggested this component enhanced attention to positive feedback as a motivationally salient stimulus (Polich, 2007; see Glazer et al., 2018 to review). Also, we did not observe any effect of addiction on this valence-dependent component which is in agreement with the previous findings in other groups of individuals with learning problems(C. di B. Luft et al., 2013) including MUD addicts (Zhao et al., 2017; Zhong et al., 2020). Accordingly, one reason for this similarity could be the addict’s ability in later cognitive processing including attention allocated to process outcome valence is not impaired (inspired from Polich, 2007). As mentioned above, the ERP studies conducted in the field of addiction especially feedback processing in MUDs are contradictory. Thereupon, future research is required to unveil the possible mechanism of these discrepancies. Taken together, the aberration in task performance, combined with intact early “FRN” and later “P300” feedback processing, may reflect latent deficiencies to assign values (weigh) to different options (stimulus)(Martin et al., 2018). This impairment (less sensitivity to value difference) may lead to difficulties in estimating response-outcome contingencies and impose a negative effect on goal-directed behavior of individuals; it may encode in an oscillatory dynamic of medial prefrontal cortex(MPFC)(Cavanagh et al., 2012; Cohen et al., 2011).

In agreement with previous studies, our time frequency decomposition results indicated that theta oscillatory activity (5–8 Hz) of MPFC (MFT) following negative feedback is associated with more enhancement than positive feedback (Cavanagh et al., 2019b; Cavanagh & Frank, 2014; C. di B. Luft et al., 2013; Van de Vijver et al., 2014). Our investigation also revealed MUDs showed a significant reduction in theta synchronization following negative feedback as compared to HCs. This hyposensitivity to unpleasant outcome in MUDs (as individuals with deficient feedback-based learning mechanisms) reconciled with previous works in other groups with feedback processing impairments including addicts(Kamarajan et al., 2012; Oberg et al., 2011) and healthy low-learners(C. di B. Luft et al., 2013). The MFT oscillations (the signature of dACC activation) are assumed to mirror the prediction error and these oscillations have an association with the option’s value assignment system(Cavanagh et al., 2019b; Cavanagh & Frank, 2014; Gläscher et al., 2009; Khamassi & Quilodran, 2010; Martin et al., 2018; Morris et al., 2011; Nieuwenhuis et al., 2004; van de Vijver et al., 2011). Therefore, punishment insensitivity in MUDs may be a result of both dopamine system dysfunction (PE modulator) and abnormal activity in areas related to options value assignment (options value sensitivity) importantly in ACC. This abnormal activity is encoded in MFT response to negative feedback. Subsequently, theta power correlation with the options value sensitivity across individuals strengthened such suggestion. Since this association has been missed in MUDs compared to HCs, it seems that addict individuals have difficulties in employing MPFC for optimal adjustment of the option’s value that is used to optimally adjust their behavior.

Furthermore, by analysis of mutual-connectivity pattern in theta range (4-6 Hz) during feedback processing we found that MPFC as a hub of performance monitoring system has communication with the RPFC (a key region in cognitive control network) (Cavanagh et al., 2009, 2010; van de Vijver et al., 2011; Watts et al., 2018). This phase locking changes, independent to the groups, enhanced more for negative feedback than positive feedback. Our results echo the finding of the previous investigation suggesting that this connectivity provide a mechanism to increase cognitive control following negative outcome to efficiently adjust subsequent behaviors. Consistent with previous intracranial electrocorticography evidence(Oehrn et al., 2014; E. H. Smith et al., 2015) our directed connectivity results estimate that this information propagation stems from the MPFC. Contrary to luft et al. we proposed that these synchronization measures are not directly correlated with the learning and the option value updating mechanism(C. di B. Luft et al., 2013). This suggestion is based on no correlation founded with behavioral performance neither at the group level nor at the individual level. Together, our second and third hypothesis partially supported. Briefly, we demonstrated that MPFC does not work properly in MUDs, which may cause insensitivity to negative feedback and lead to non-optimal assignment value to options. Notably, the connectivity of this monitoring-related area with the cognitive control-related remains intact in MUDs. Although such findings may seem contradictory at the first glance and raise the question of whether the MPFC is malfunctioning, the abovementioned synchrony should also be damaged. But it is worth considering that the MPFC-RPFC connectivity may be due to the third subcortical area, which is suggested to be taken into consideration for future investigations.

Our investigation of brain-behavior linkage echoes the finding of Cavanagh and colleagues (Cavanagh et al., 2011) who demonstrated that coupling between error signal (the MFT power) and avoidance learning (Nogo learning accuracy in the test phase) was marginally enhanced in the healthy group. This normative association appears to be highly blunted in the addict group, despite the absence of differences in test-phase behavioral performance with HCs (supporting fourth hypothesis). Together, our results suggest that the response-outcome association system reflected in MFT, defecated in MUDs. In other words, this altered mechanism could be related to the effects of inattention/insensitivity to negative valance-feedback in addicts. If this be the case, it means that addicts learn less from negative feedback and have smaller negative prediction error, which shows itself in decreased error signal–punishment learning coupling. Again, a mechanistic explanation of this effect may be downregulation of D2 receptors in the addict’s midbrain due to chronic methamphetamine use(Groman et al., 2018; Harlé et al., 2015; Lim et al., 2021). This role, error signal-punishment learning interaction, is also supported by the neural network model of cortico-striatal(Frank & O’Reilly, 2006).

## CONCLUSIONS AND FUTURE DIRECTIONS

Many emerging studies in the field of cognitive psychology have pointed out the importance of brain mechanism as the method of diagnosis and classifying diseases(Cavanagh et al., 2019b; Kirschner & Klein, 2022b; McLoughlin et al., 2022). Therefore, relying on the unique ability of EEG to may reveal potential biomarkers, in this study we have investigated the brain feedback-related mechanisms in addicted individuals. Consequently, we examined the separable effects of methamphetamine consumption in four domains: (a) behavioral performance, (b) performance of the related computational model, (c) electrocortical responses correlate of behavior, and (d) behavioral performance-frequency association. Our analysis revealed, methamphetamine abusers did not differ in performance measures of learning (a) while they are less sensitive to value difference (b). We demonstrated MUDs hyposensitivity to negative feedback which is reflected in medial-frontal theta-band power (c) plays important role in this deficiency. Following this assumption, the missed correlation (d) between the value sensitivity and mentioned oscillatory activity in MUDs estimated that probably this lack due to theta power alone and not any communication between PFC feedback processing-related areas. In sum, we suggested a responsible mechanism for assigning value to the options, reflected in MFT, is impaired in addicts. This malfunction, which shows itself at a behavioral level in both smaller negative prediction error and especially lower in option value sensitivity, is complying with the behavior of addicts in the real world: they are insensitive to negative consequences of drug use. According to connectivity analysis, we also suggested that the dynamic cycle that deals with monitoring behavior and subsequently increasing control after negative feedback is intact in addicts. Finally, we conclude that the computational model in combination with neurophysiological correlated responsible for negative feedback including ERPs, connectivity between task-related areas, and MFT can reveal different aspects of learning from feedback in these individuals subject to dopamine alteration.

In addition to MFT being a reflection of both PE and options value sensitivity it is also related to associative and working memory(Backus et al., 2016). Therefore, considering these functions that are involved in the valuation system leads to a more sophisticated understanding of the altered feedback processing mechanisms in addicts (see Yoo & Collins, 2022 to inspiration and overview). Since the cause of the behavior-brain relationship may be more the phase of theta oscillations than its amplitude, a more detailed examination of the phase association instead of the amplitude in MPFC is suggested (see Duprez et al., 2020 for inspiration). On the other hand, investigating the effect of regulating theta oscillations as a therapeutic approach can open new windows in the treatment of the addicts (similar to Grover et al., 2021). Finally, this is the remained gap to be filled by future researches: impairment we observed in our study related to feedback system dysfunction or general neurocognitive dysfunction for performing any given cognitive task in addicts?

