## supplementary material for "Dysfunctional feedback processing in methamphetamine abuser; evidence from neurophysiological and computational analysis"

**A:** *Q***-Learning with separate learning parameter**

We implemented the *Q*-learning model in *PyStan* 2.19 (Carpenter et al., 2017) using the Markov chain Monte Carlo (MCMC) algorithm(Gelman et al., 1996). In addition, we considered iteration and chains values of 8000 and 4, respectively, and *warmup* is set equal to 40% of iterations, ergo the samples are obtained by guaranteeing the convergence condition$\hat{R}<1.01$. For this purpose we defined priors/hyper-priors distribution as follows:

$$\mu_{\eta_{\left( +/- \right)}}\mathcal{\sim N}\left( 0, 1 \right);\sigma_{\eta_{\left( +/- \right)}}\mathcal{\sim HN}\left( 0.3 \right), \eta_{\left( +/- \right)}\sim\phi(\mathcal{N(}\mu_{\eta_{\left( +/- \right)}},\sigma_{\eta_{\left( +/- \right)}}))$$

$$\mu_{s}\mathcal{\sim N}\left( 1,10 \right);\sigma_{s}\mathcal{\sim HN}\left( 2 \right);s\sim\mathcal{N(}\mu_{s}, \sigma_{s})$$

Where $\mathcal{N, HN}$ and $\phi$ indicate normal distribution, half normal distribution and normal CDF, respectively.

We compared two *Q*-RL models on 17 HC and 14 MUD data for the learning process, one with one learning parameter and the other with separating positive and negative learning parameters. Among the models presented for both groups, the better models in predicting participants' behavior include separate learning rates (η) to fit positive and negative prediction errors. In the quantitative process of model comparison, a model that has a lower value of WAIC and a higher value of lppd is more acceptable, which in table 1 clearly shows the better results of the second model for these two values compared to the first model (For both groups). Next, the posterior predictive checks (PPC), as qualitative model comparison approach, conducted. Briefly PPC estimated for mean accuracy and mean RT searched whether the 95% BCIs of the posterior predictive distributions contained the participants mean accuracy and RTs. The PPC results of model 2 given in the following show that model with separate learning rate has well predicted the behavior of the learning in both groups (Fig. 9). Taken together, posterior of group mean parameters distributions for healthy control (HC) and methamphetamine use dependent group as well as the contrast for HC-MA are illustrated in table 2.

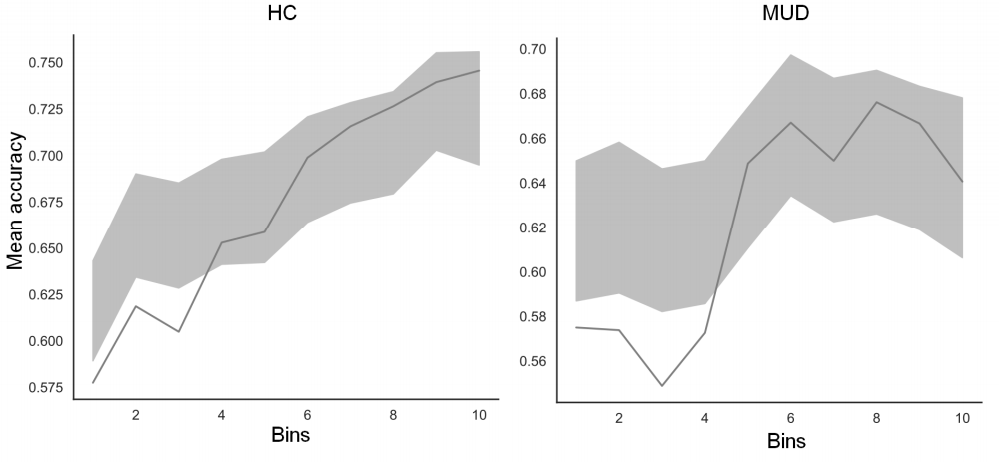

Fig 1| Posterior predictive checks of mean accuracy according to the RL with separated learning rate for HC (left) and MUD (right) groups. The 95% Bayesian credible interval (BCI) of the posterior predictive distribution (*shaded areas*) and the observed accuracy (solid line) for bins of trials within the PLT training phase is illustrated.

Table 1| Quantitative model comparison between the reinforcement learning models.

| Model | $\boldsymbol{\eta}$ | $\boldsymbol{sen}$ | HC  WAIC | *p*WAIC | -lppd | Rank | MUD  WAIC | *p*WAIC | −lppd | Rank |
| --- | --- | --- | --- | --- | --- | --- | --- | --- | --- | --- |
| 1 | Fixed | Fixed | 11694 | 76 | 5810 | 2 | 10540 | 53 | 5250 | 2 |
| 2 | Separated | Fixed | 11124 | 84 | 5528 | 1* | 10111 | 63 | 5031 | 1* |

Table 2| Between group posterior parameters distributions of the RL model with separated learning rate results. HDI represent the 95% highest density intervals, BF represent Bayes factor for directional effects. BF>1 indicated reduction in a parameter for MUD vs. HC and vice versa for BF<1.

|  | HC | | | MUD | | | HC-MUD | | | |
| --- | --- | --- | --- | --- | --- | --- | --- | --- | --- | --- |
|  | m | HDI |  | m | HDI |  | m | HDI |  | BF |
| $\boldsymbol{\eta}_{\boldsymbol{+}}$ | 0.0438 | 0.0254 | 0.0679 | 0.0416 | 0.0244 | 0.0634 | 0.0022 | 0.0254- | 0.0325 | 1.4199 |
| $\boldsymbol{\eta}_{\boldsymbol{-}}$ | 0.0007 | 0.0000 | 0.0020 | 0.0002 | 0.0000 | 0.0010 | 0.0004 | 0.0005- | 0.0017 | 3.5124 |
| $\boldsymbol{sens}$ | 3.2980 | 2.3449 | 4.4037 | 2.3562 | 1.4961 | 3.2667 | 0.9418 | -0.3502 | 2.3425 | 10.130 |

**B: Bayesian linear and logistic regression**

For regression models, ten levels of blocks are used to build predictors matrix to test their main effect on accuracy (coded logically) and RTs for separate conditions. Block was coded as 1 to 10 respectively.

In sum the implementation of the model is similar to the learning model in Supplementary material A. For the regression analysis of RT, we applied the Bayesian linear regression model, in which the priors, hyper-priors, and the model are specified as follows:

$$\lambda_{1}\mathcal{\sim N}\left( 0, 2 \right);\lambda_{2}\mathcal{\sim N}\left( 0, 2 \right)$$

$$\varepsilon\mathcal{\sim HN}\left( 1 \right)$$

$$\bar{RT}_{t}=\lambda_{1}+M_{t}\lambda_{2}$$

$$RT_{t}\mathcal{\sim N}\left( \bar{RT}_{t},\varepsilon\right)$$

here$\lambda_{1}$, $\lambda_{2}, M$ and $\varepsilon$ represent the intercept vector, coefficient vector, prediction matrix and noise term, respectively. In addition, $\mathcal{N}$indicate normal distributions parameterized with mean and standard deviation and $\mathcal{HN}$ indicate half normal distribution parameterized with standard deviation.

In almost the same way, we used the Bayesian logistic regression model for regression analysis of accuracy, in which priors and hyper-priors are similar to the linear model and the only difference with the linear model is the choice of the Bernoulli distribution in the prediction process as follows:

$$\bar{ACC}_{t}=logit\left( \lambda_{1}+M_{t}\lambda_{2} \right)$$

$$ACC_{t}\sim Bern(\bar{ACC}_{t})$$
